# Dim light at night impacts circadian rhythms and Alzheimer’s disease-like neuroinflammation and neuropathology in humanized APP SAA knock-in mice

**DOI:** 10.1101/2025.10.02.680043

**Authors:** Marilyn J. Duncan, Margaret R. Hawkins, Leke Bytyqi, Haleigh R. Whitlock, Savannah Shepard, MaKayla F. Cox, Esther G. Drinkard, Teresa Macheda, Kelly N. Roberts, Katherina Kohler, Mary-Claire Schmidt, Carrie E. Johnson, Sridhar Sunderam, Bruce F. O’Hara, Michael P. Murphy, Adam D. Bachstetter

## Abstract

Artificial light at night (light pollution) is widespread but understudied in the context of Alzheimer’s disease (AD). Sleep and circadian disruption have been linked to amyloid-β (Aβ) accumulation and neuroinflammation, but whether dim light at night (dLAN) modifies these processes remains unclear. We tested whether chronic dLAN exposure (8 lux during the dark phase, 8 weeks) alters circadian rhythms, amyloid pathology, and neuroinflammation in 12–13 month-old humanized APP knock-in (KI) mice. hAPP^SAA^ KI mice, which develop plaques, were compared with hAPP^WT^ KI controls carrying only a humanized APP sequence. dLAN reduced circadian rhythm amplitude and stability while increasing fragmentation in both genotypes within two weeks. In hAPP^SAA^ KI mice, dLAN modestly increased hippocampal plaque burden and soluble neocortical Aβ. Astrocyte reactivity was elevated by genotype but not altered by nighttime light exposure. In contrast, microglial markers (CD45, MHCII) were increased with dLAN with CD45+ area elevated in hippocampus, and MHCII+ cell counts greater in the cortex and hippocampus of hAPP^SAA^ KI mice. There were also distinct spatial responses between the microglia markers suggesting that dLAN primes microglia toward an antigen-presenting phenotype (MHCII) in the presence of Aβ. Yet, the microglia/macrophage priming was not associated with amplified cytokine or chemokine levels at the 8-week dLAN exposure timepoint in the brain. These findings add to growing evidence that nighttime light exposure can disrupt circadian and immune regulation, and suggest that environmental light pollution should be further explored as a modifiable factor contributing to Alzheimer’s disease progression.

**Statement of Significance:** Light at night is a common feature of modern life, yet its influence on Alzheimer’s disease remains poorly understood. We show that dim light at night disrupts circadian rhythms, modestly increases amyloid pathology, and shifts microglia toward an antigen-presenting state in an amyloid-prone model. These findings identify light at night as a modifiable factor that may worsen risk or progression of neurodegenerative disease. A critical gap is whether circadian and immune changes resolve after darkness at night is restored. If they persist, early exposure could leave lasting imprints on brain aging. Addressing this question is essential for guiding strategies to mitigate the impact of light pollution.

## Introduction

Disruptions of circadian rhythms and sleep are associated with increased risk of Alzheimer’s disease and related dementias (ADRD)^1–3^. Among the many factors that disturb circadian timing, exposure to artificial light at night, commonly referred to as light pollution, is widespread in modern environments ^4^. Light at night comes from both indoor sources such as lamps and electronic devices and outdoor sources including streetlights and billboards. Even low-intensity light at night (5–10 lux) has been shown to disrupt sleep in humans, reducing total sleep time, increasing wakefulness after sleep onset, and lowering sleep efficiency ^5^ ^6,7^. Nighttime light exposure above 10 lux has also been linked with greater sleep fragmentation ^8^. Sleep fragmentation itself is a risk factor for Alzheimer’s disease (AD) as well as a factor that predicts conversion from mild cognitive impairment (MCI) to AD and acceleration of AD progression ^9–12^.

Experimental studies in mice have shown that sleep fragmentation can drive AD-related pathology, including accumulation of amyloid-β (Aβ) and neuroinflammation mediated by activated microglia ^13^. In parallel, several groups have reported that dim light at night exposure increases hippocampal microglial activation and inducible Nitric Oxide Synthase (iNOS, *an enzyme that produces nitric oxide*) expression even without immune challenge, and amplifies CNS cytokine responses to lipopolysaccharide, high-fat diet, or particulate matter ^14,15 16^. Together, these findings suggest that circadian disruption by light at night has the capacity to engage inflammatory pathways in the brain.

To investigate how these effects intersect with amyloid pathology, we used two humanized APP knock-in mouse lines that differ in their propensity to develop Aβ deposition. The hAPP^WT^ KI line carries a humanized Aβ sequence but does not develop plaques, whereas the hAPP^SAA^ KI line carries additional familial AD-linked mutations that promote amyloidogenic processing and robust plaque deposition with age ^17–19^. We hypothesized that dim light at night would exacerbate amyloid accumulation and microglial activation in this sensitized background. To test this, we measured circadian activity rhythms, amyloid pathology, and glial responses in both knock-in lines after chronic light exposure.

## Methods

### Animals and housing

All animal procedures were approved by the Institutional Animal Care and Use Committee of the University of Kentucky and were conducted in accordance with the NIH Guide for the Care and Use of Laboratory Animals (Protocol: 2018-3066). A total of 39 mice aged 12–13 months were studied: hAPP^SAA^ KI mice (RRID:IMSR_JAX:034711; N = 20; 10F/10M) and hAPP^WT^ KI controls (RRID:IMSR_JAX:033013; N = 19; 10F/9M). The hAPP^SAA^ KI strain carries a humanized Aβ region (G676R, F681Y, R684H) together with the Swedish (KM670/671NL), Arctic (E693G), and Austrian (T714I) mutations, resulting in progressive plaque deposition and microgliosis in vivo without altering endogenous gene regulation ^17^. The hAPP^WT^ KI strain contains the humanized Aβ sequence but no familial AD mutations. Mice were singly housed in light-tight, sound-attenuated cabinets with ventilated racks and ad libitum access to food (Teklad Global 18% Protein Rodent Diet, 2918) and water. A 12:12 light–dark cycle (200 lux white light during the light phase, complete darkness during the dark phase) was maintained for one week of baseline recording. Mice were then randomly assigned within genotype and sex to remain on this standard cycle (LD) or to receive dim light at night (dLAN), in which the dark phase was replaced with 8 lux white light (measured with Gigahertz Optic MSC15). Final group sizes were: hAPP^WT^ KI-LD (n = 7, 4F/3M), hAPP^WT^ KI-dLAN (n = 12, 6F/6M), hAPP^SAA^ KI-LD (n = 12, 7F/5M), and hAPP^SAA^ KI-dLAN (n = 8, 4F/4M).

### Circadian activity monitoring

Circadian rest-activity rhythms were monitored with passive infrared (PIR) motion detectors (ClockLab Wireless PIR Nodes, Actimetrics v6.1). Because only 20 sensors were available, recordings were collected in alternating weeks, with each cohort monitored on odd-numbered weeks throughout the study. Cosinor analysis fitted a standard sine wave to the data, which was used to determine circadian rhythm parameters including: amplitude (distance from the trough to the peak of the sine wave and a measure of rhythm robustness), acrophase (time of the peak of the rhythm), and the midline estimating statistic of rhythm (MESOR, the mean of the model). Non-parametric circadian rhythm analysis was used to determine two additional circadian parameters: intradaily variability (IV, a measure of rhythm fragmentation, range 0-3) and interdaily stability (IS, a measure of rhythm stability from day-to-day, range 0-3) ^20^. Dark- and light-phase activity counts were separately quantified to assess phase-specific changes.

### Tissue collection and processing

Mice were euthanized at Zeitgeber Time (ZT) 7–8 by CO₂ asphyxiation followed by decapitation, after completing eight weeks of light treatment. Brains were rapidly removed and bisected sagittally. One hemibrain was drop-fixed in 4% paraformaldehyde overnight, cryoprotected in 30% sucrose/PBS, and coronally sectioned at 30 μm on a freezing microtome. Sections were stored in cryoprotectant at -20 °C. The other hemibrain was microdissected into neocortex and hippocampus, flash-frozen in liquid nitrogen, and stored at -80 °C for biochemical assays.

### Immunohistochemistry

Systematic uniform random sampling was implemented by selecting every 10th 30-µm coronal section across 1.3–2.5 mm posterior to bregma, resulting in approximately six sections per region per mouse. Free-floating sections were immunostained as described previously ^13,21,22^. Primary antibodies included: rabbit anti-GFAP (1:10,000, Dako Z0334, RRID:AB_10013382), rat anti-CD45 (1:1,000, BioLegend 103102, RRID:AB_312967), rat anti-MHCII (I-A/I-E; 1:1,000, BioLegend 107601, RRID:AB_313317), and mouse anti-Aβ (clone 6E10, 1:3,000, BioLegend 803007, RRID:AB_2564657). Sections were incubated overnight at 4 °C, followed by biotinylated secondary antibodies and detection with DAB (Vector Laboratories). Slides were scanned on a Zeiss AxioScan Z.1, and regions of interest (neocortex, hippocampus) were manually outlined. Positive-staining area (% area fraction) was quantified in HALO-AI (Version 4.1, Indica Labs) using a single, fixed threshold per marker applied uniformly across all slides. For object counts (CD45⁺ and MHCII⁺ clusters), a custom-trained HALO AI classifier was used to segment immunoreactive objects; the model was trained on manually annotated tiles spanning genotypes and light conditions and locked prior to batch analysis.

Segmented objects were exported with area (µm²) and centroid coordinates, normalized as counts per mm², and binned by size using quartile-derived cut points (small <216 µm², medium 216–956.5 µm², large >956.5 µm²). All analyses were performed blind to genotype and treatment, and markup overlays were reviewed for quality control.

### Tissue extraction for biochemical assays

As previously described ^23^, frozen neocortical and hippocampus tissues were homogenized on ice using an Omni Bead Ruptor (Omni International) in PBS-based lysis buffer (1:20 w/v) containing 1 mM EDTA and protease inhibitors (1 mM PMSF, 1 µg/ml leupeptin). Samples were centrifuged at 12,000 × g for 20 min at 4 °C, and the supernatant was collected as the PBS-soluble fraction. The pellet was rehomogenized in detergent-containing buffer (T-PER, ThermoFisher #78510, supplemented with Halt Protease and Phosphatase Inhibitor Cocktail, ThermoFisher #78442) and centrifuged as above; the supernatant was collected as the detergent-soluble fraction. Finally, the remaining pellet was solubilized in 70% formic acid, centrifuged at 12,000 × g for 20 min. Aliquots of all fractions were stored at –80 °C until analysis. Protein concentration in each fraction was determined using the bicinchoninic acid (BCA) assay (Thermo Scientific).

### Aβ extraction and ELISA

As previously described ^24^, sandwich ELISAs were performed on Immunolon 4HBX plates (Thermo Fisher), coated with human-specific Aβ1–16 antibody 42.5 (1 µg/well), and detected with biotinylated anti-Aβ17–24 (4G8, BioLegend 800701, RRID:AB_2728526) and Neutravidin-HRP (Thermo Fisher 31001). FA samples were neutralized via a 1:20 dilution in 1.0 M Tris-Base / 0.5 M Na2HPO4 and diluted 1/800 prior to use and immediately after thawing. Recombinant Aβ42 (rPeptide A-1163-2) was used for standards. Absorbance was measured at 450 nm (BioTek plate reader), and concentrations were interpolated using standard curves. All values were normalized to total protein content (BCA assay, Thermo Fisher 23227).

### Cytokine and chemokine quantification

PBS-soluble fractions from neocortex and hippocampus were assayed using custom Meso Scale Discovery (MSD) V-PLEX panels (K15245D, K15048D) per manufacturer’s instructions, as previously described ^23^. Analytes included IL-1β, IL-6, IL-10, TNFα, IL-17A, CCL2 (MCP-1), CCL3 (MIP-1α), CXCL1 (KC/GRO), CXCL2 (MIP-2), and CXCL10 (IP-10). Samples (50 μL, 100–200 μg protein) were incubated overnight at 4 °C in pre-coated plates. Plates were washed with the BioTek 50TS plate washer, probed with detection antibodies, and read on a MESO QuickPlex SQ 120. Concentrations were interpolated from 8-point standard curves. All values were normalized to total protein content (BCA assay, Thermo Fisher 23227).

### Statistical analysis

All analyses were performed using JMP Pro 17 (SAS Institute, Cary, NC), and graphs were generated with GraphPad Prism version 10.4.2 (GraphPad Software, San Diego, CA). Animals were assigned to treatment groups using randomization procedures, with the constraint that littermates were distributed across conditions to avoid confounding by family effects. Because of animal availability, group sizes were not fully balanced. To reduce temporal bias, animals were run in two cohorts of ∼20 mice, with randomized assignment to ensure approximately equal representation of genotypes and light conditions within each cohort. For histological and biochemical assays, samples from different groups were randomized and interleaved across staining batches and assay plates to minimize batch effects. Each animal was assigned a unique identifier that concealed genotype and treatment. All experiments, tissue processing, and image analyses were performed blind to group, and codes were broken only after data collection was complete; statistical analyses were then conducted by ADB. Sex was included as a factor in initial models, but no main effects or interactions were detected; data were therefore pooled across sexes. Circadian rest–activity rhythms were analyzed using mixed models for repeated measures, with light condition, genotype, and week of treatment as fixed effects and mouse ID as a random effect. Histological and biochemical endpoints were analyzed with two-way ANOVA (genotype x light), followed by planned within-genotype contrasts. Model assumptions were evaluated using the Shapiro–Wilk test for normality and Levene’s test for homogeneity of variance. When assumptions were violated, Box-Cox transformations were applied. Effect sizes were reported as partial η². Significance was set at p < 0.05. Data are presented as mean ± SEM, with individual animal values shown in figures. Source data and a data dictionary have been deposited in Dryad and will be publicly available upon publication (DOI: [pending]).

## Results

### Dim light at night impairs circadian activity patterns in both hAPP^WT^ and hAPP^SAA^ mice

Circadian activity rhythms were quantified to determine whether dim light at night alters the amplitude and stability of the daily rest–activity cycle (**Fig. 1**). Exposure to dim light at night reduced rhythm amplitude, indicating a less robust distinction between active and rest phases (**Fig. 1A**; F(1,35) = 7.89, p = 0.008), and lowered the mean 24-h activity level (MESOR), reflecting a decrease in overall daily activity level (**Fig. 1B**; F(1,35) = 6.04, p = 0.019). Dim light at night increased intradaily variability (IV), reflecting greater fragmentation of activity within each 24-h cycle (**Fig. 1C**; F(1,35) = 7.99, p = 0.008), and reduced interdaily stability (IS), indicating less consistent day-to-day timing of activity (**Fig. 1D**; F(1,35) = 32.36, p < 0.0001). To determine whether reduced amplitude reflected changes during specific phases of the light–dark cycle, we analyzed dark- and light-phase activity separately. Dim light at night suppressed activity during the dark (active) phase (**Fig. 1E**; F(1,35) = 8.97, p = 0.005), while light-phase activity remained unchanged (**Fig. 1F**; F(1,35) = 0.16, p = 0.69). Disruptions in circadian activity rhythms were evident by the second week of exposure and remained throughout the recording period, as indicated by significant main effects of treatment, and by treatment × week interactions for IV (**Fig. 1C**; F(4,140) = 3.31, p = 0.013), IS (**Fig. 1D**; F(4,140) = 7.99, p = 0.008) and dark (active) phase (**Fig. 1E**; F(4,140) = 4.80, p = 0.001). Post hoc contrasts indicated the largest differences at week 2 (p < 0.01), with attenuation of effects by week 8. There were no main effects of genotype or genotype × light interactions (all p > 0.15), showing that dim light at night impaired rhythms similarly in hAPP^WT^ KI and hAPP^SAA^ KI mice. These findings demonstrate that dim light at night broadly disrupts circadian organization in both hAPP^WT^ KI and hAPP^SAA^ KI mice, supporting its use as a model to investigate how circadian disruption contributes to Alzheimer’s disease–related neuropathological changes.

**Figure 1.**
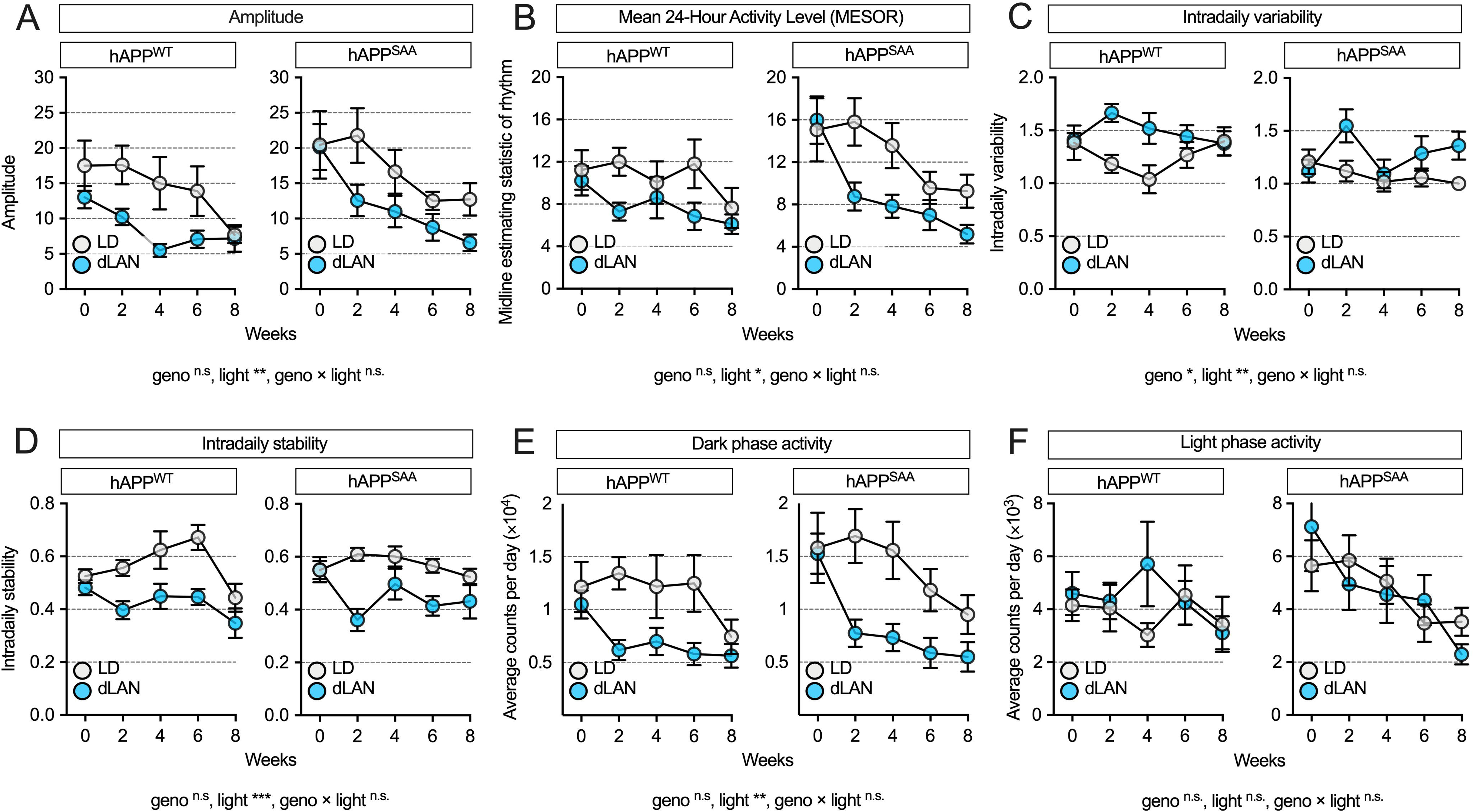
Chronic exposure to dim light at night (dLAN) disrupts circadian activity rhythms. **(A)** Amplitude of the daily activity rhythm was reduced by dLAN across both hAPP^WT^ KI and hAPP^SAA^ KI mice.**(B)** The MESOR, reflecting the mean 24-h activity level, was also decreased by dLAN. (C) Intradaily variability (IV), a measure of rhythm fragmentation, was increased by dLAN, with an additional main effect of genotype. **(D)** Interdaily stability (IS), indicating day-to-day regularity of activity, was reduced by dLAN. **(E)** Dark-phase activity counts were decreased by dLAN, whereas **(F)** light-phase activity counts were unaffected. Data represent group means ± SEM across 8 weeks of exposure. Sample sizes were: hAPP^WT^ KI-standard 12/12 light–dark cycle (LD) (n = 7, 4F/3M), hAPP^WT^ KI-dLAN (n = 12, 6F/6M), hAPP^SAA^ KI LD (n = 12, 7F/5M), hAPP^SAA^ KI -dLAN (n = 8, 4F/4M). Statistical analyses were conducted using linear mixed-effects models with lighting condition (LD vs dLAN), genotype (hAPP^WT^ KI vs hAPP^SAA^ KI), sex, and week of treatment as fixed effects, and mouse ID nested within genotype and treatment as a random effect. Main effects and interactions are reported below each panel; symbols denote significance levels (*p<0.05, **p<0.01, ***p<0.001).

### Dim light at night modestly increases Aβ in hAPP^SAA^ KI mice

Immunohistochemistry for Aβ (6E10) revealed abundant plaques in neocortex and hippocampus of hAPP^SAA^ KI mice, with no detectable plaques in hAPP^WT^ KI controls (**Fig. 2A-C**). Genotype strongly affected neocortical plaque burden (6E10+ area) (F(1,35) = 97.37, p < 0.001, partial η² = 0.736), whereas light condition and the genotype × light interaction were not significant (**Fig. 2D**). Plaque-size distributions in neocortex showed robust genotype effects across small, medium, and large plaques (all p < 0.001; **Fig. 2E**).

**Figure 2.**
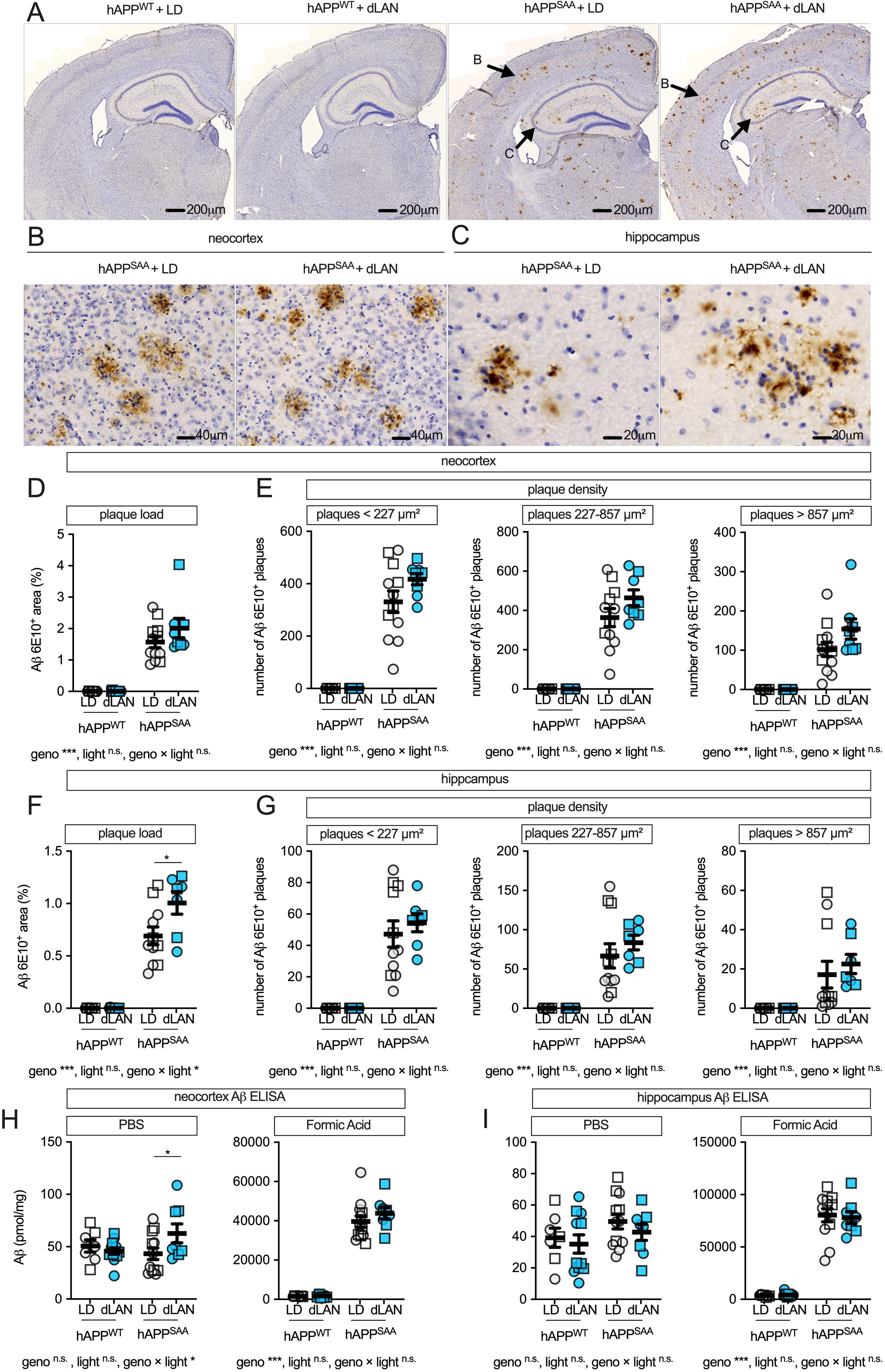
Effect of dim light at night on neocortical and hippocampal Aβ pathology. **(A)** Low-magnification 6E10 immunostaining in coronal sections from hAPP^WT^ KI and hAPP^SAA^ KI mice maintained under a standard 12/12 light–dark cycle (LD) or dim light at night (dLAN). **(B)** Higher magnification of neocortex and **(C)** hippocampus illustrating Aβ plaques (brown DAB; hematoxylin counterstain). Scale bars: A, 200 µm; B, 40 µm; C, 20 µm. **(D)** Quantification of neocortical Aβ 6E10+ area. **(E)** Neocortical plaque density stratified by size: <227 µm² (small), 227–857 µm² (medium), >857 µm² (large). **(F)** Hippocampal Aβ 6E10+ area. **(G)** Hippocampal plaque density stratified by size bins as in **(E)**. ELISA of Aβ from neocortex **(H)** and hippocampus **(I)**, extracted with PBS (soluble) or formic acid (insoluble); values expressed as fmol/mg protein. Symbols represent individual mice; circles denote females and squares denote males; bars show mean ± SEM. Sample sizes were: hAPP^WT^ KI-LD (n = 7, 4F/3M), hAPP^WT^ KI-dLAN (n = 12, 6F/6M), hAPP^SAA^ KI-LD (n = 12, 7F/5M), hAPP^SAA^ KI -dLAN (n = 8, 4F/4M). Analyses were conducted using two-way ANOVA (genotype × light) with planned within-genotype contrasts where appropriate. Main effects and interactions are indicated below the panels. Asterisks denote significance levels (*p < 0.05, **p < 0.01, ***p < 0.001).

In hippocampus, plaque burden was increased by genotype (F(1, 33) = 163.04, p < .001, partial η² = 0.832), and genotype × light interaction was significant (F(1,33) = 5.61, p = 0.024, partial η² = 0.145; **Fig. 2F**). Post hoc contrasts confirmed greater hippocampal Aβ area in hAPP^SAA^ KI mice under dim light at night compared with the standard light–dark cycle (p = 0.0048), with no effect in hAPP^WT^ KI (p = 0.45). Hippocampal plaque-size distributions were increased by genotype (all p < 0.001), with no effects of light or interaction (**Fig. 2G**).

Biochemical Aβ measurements showed a regionally distinct pattern of Aβ changes compared to histology. In neocortex, PBS-soluble Aβ showed a genotype × light interaction (F(1, 35) = 4.22, p = 0.047, partial η² = 0.108). Pot hoc analysis showed that hAPP^SAA^ KI mice exposed to dim light at night had higher soluble Aβ than those in the standard light–dark cycle (p = 0.0256), whereas hAPP^WT^ KI showed no light effect (**Fig. 2H**). Neocortical formic acid-soluble (insoluble) Aβ was elevated by genotype (F(1,35) = 325.28, p < 0.001) and was unaffected by light condition or interaction (**Fig. 2H**). In hippocampus, PBS-soluble Aβ did not differ by genotype or light, whereas formic acid–soluble Aβ was elevated in hAPP^SAA^ KI mice (F(1,34) = 249.99, p < 0.001) and was not altered by light (p = 0.87) or interaction (p = 0.76; **Fig. 2I**). Thus, dim light at night produced a modest, region-specific increase in Aβ pathology, most evident as greater hippocampal plaque burden and elevated neocortical soluble Aβ in hAPP^SAA^ KI mice.

### Astrocyte reactivity is elevated in hAPP^SAA^ KI mice and is not altered by dim light at night

GFAP immunohistochemistry revealed sparse labeling in hAPP^WT^ KI mice and robust astrocytic staining in hAPP^SAA^ KI mice across neocortex and hippocampus (**Fig. 3A-L**). Quantitatively, neocortical GFAP+ area was strongly increased by genotype (F(1,34) = 333.57, p < 0.001, partial η² = 0.908) with no effect of light condition and no genotype × light interaction (**Fig. 3M**). In hippocampus, GFAP+ area was also elevated by genotype (F(1,34) = 8.08, p = 0.008, partial η² = 0.192), and the interaction was not significant (**Fig. 3N**). Thus, astrocyte reactivity was markedly increased in the amyloid-prone hAPP^SAA^ KI background, and dim light at night did not further alter GFAP immunoreactivity at the time point studied. The large increase associated with genotype may have created a ceiling effect that would need to be explored further in future studies.

**Figure 3.**
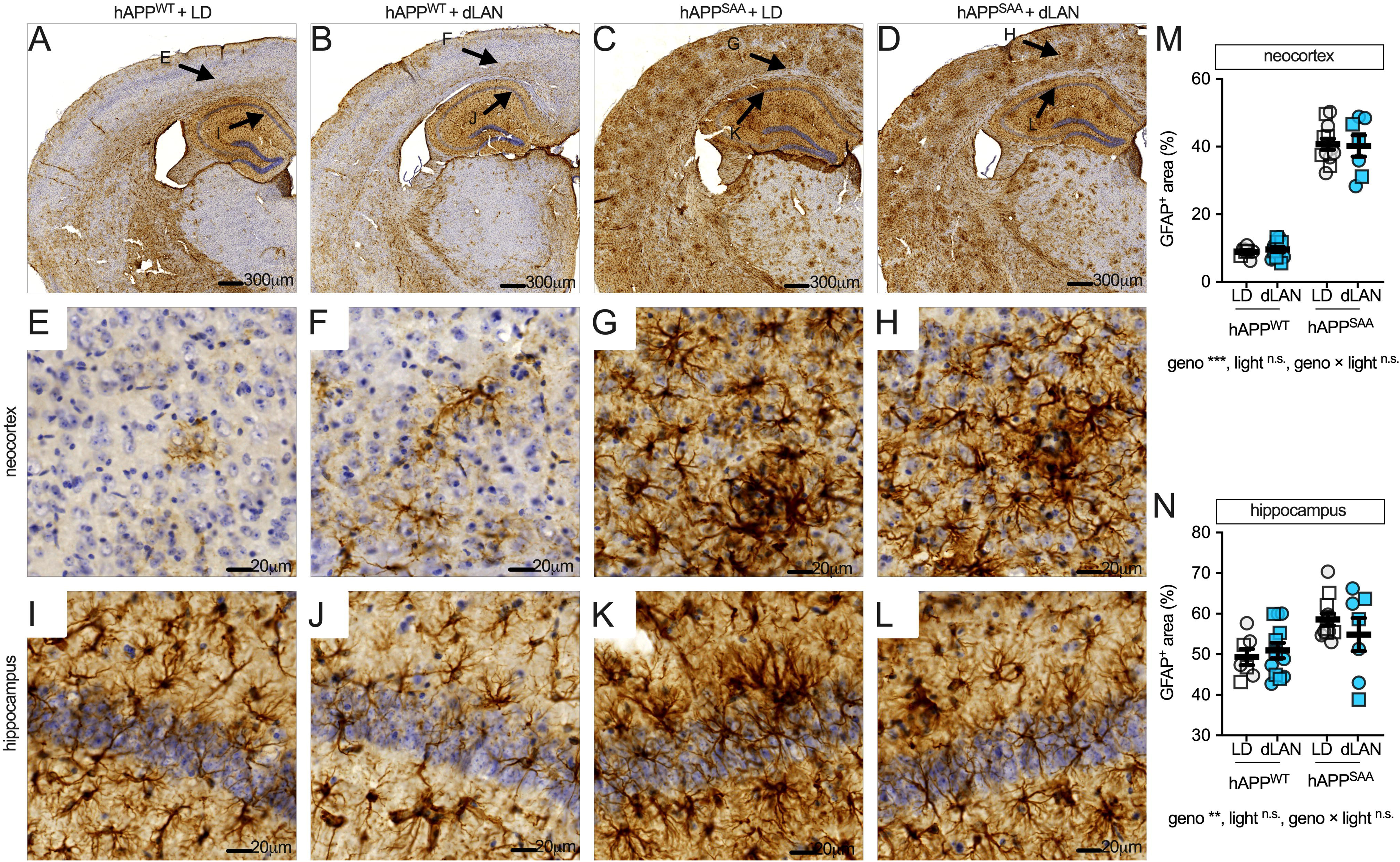
GFAP immunoreactivity is increased by amyloid genotype, with no additional effect of dim light at night detected. (A-D) Low-magnification coronal sections showing GFAP staining in hAPP^WT^ KI and hAPP^SAA^ KI mice maintained under a standard 12/12 light–dark cycle (LD) or dim light at night (dLAN). **(E-H)** Higher-magnification fields from neocortex; **(I–L)** hippocampus. Brown DAB, hematoxylin counterstain. Scale bars: A–D, 300 µm; E–L, 20 µm. **(M)** Quantification of GFAP+ area (%) in neocortex. **(N)** GFAP+ area (%) in hippocampus. Symbols represent individual mice; circles denote females and squares denote males; bars show mean ± SEM. Sample sizes were: hAPP^WT^ KI-LD (n = 7, 4F/3M), hAPP^WT^ KI-dLAN (n = 12, 6F/6M), hAPP^SAA^ KI -LD (n = 12, 7F/5M), hAPP^SAA^ KI-dLAN (n = 8, 4F/4M). Analyses were conducted using two-way ANOVA (genotype × light) with planned within-genotype contrasts where appropriate. Main effects and interactions are indicated below the panels. Asterisks denote significance levels (*p < 0.05, **p < 0.01, ***p < 0.001).

### CD45^+^ cell burden is elevated in hAPP^SAA^ KI mice and further increased by dim light at night

CD45, the leukocyte common antigen, is upregulated on reactive microglia and highly expressed on monocyte-derived macrophages. CD45 can also be detected on lymphocytes, although these were not commonly observed in our samples. Immunohistochemistry showed minimal CD45 labeling in hAPP^WT^ KI mice and dense periplaque CD45+ clusters in hAPP^SAA^ KI mice across neocortex and hippocampus with the morphological appearance of microglia (**Fig. 4A-L**). In neocortex, CD45+ area (%) was strongly increased by genotype (F(1,31) = 459.29, p < 0.001, partial η² = 0.937), with no effect of light or genotype × light interaction (both p ≥ 0.29; **Fig. 4M**). In hippocampus, CD45+ area was elevated by genotype (F(1,31) = 283.40, p < 0.001, partial η² = 0.901), and both light (F(1,31) = 5.90, p = 0.021, partial η² = 0.160) and the genotype × light interaction were significant (F(1,31) = 5.25, p = 0.029, partial η² = 0.145; **Fig. 4N**). Within-genotype contrasts confirmed higher hippocampal CD45+ area in hAPP^SAA^ KI mice exposed to dim light at night compared with the standard light– dark cycle (p = 0.0073), with no effect in hAPP^WT^ KI mice (p = 0.473).

**Figure 4.**
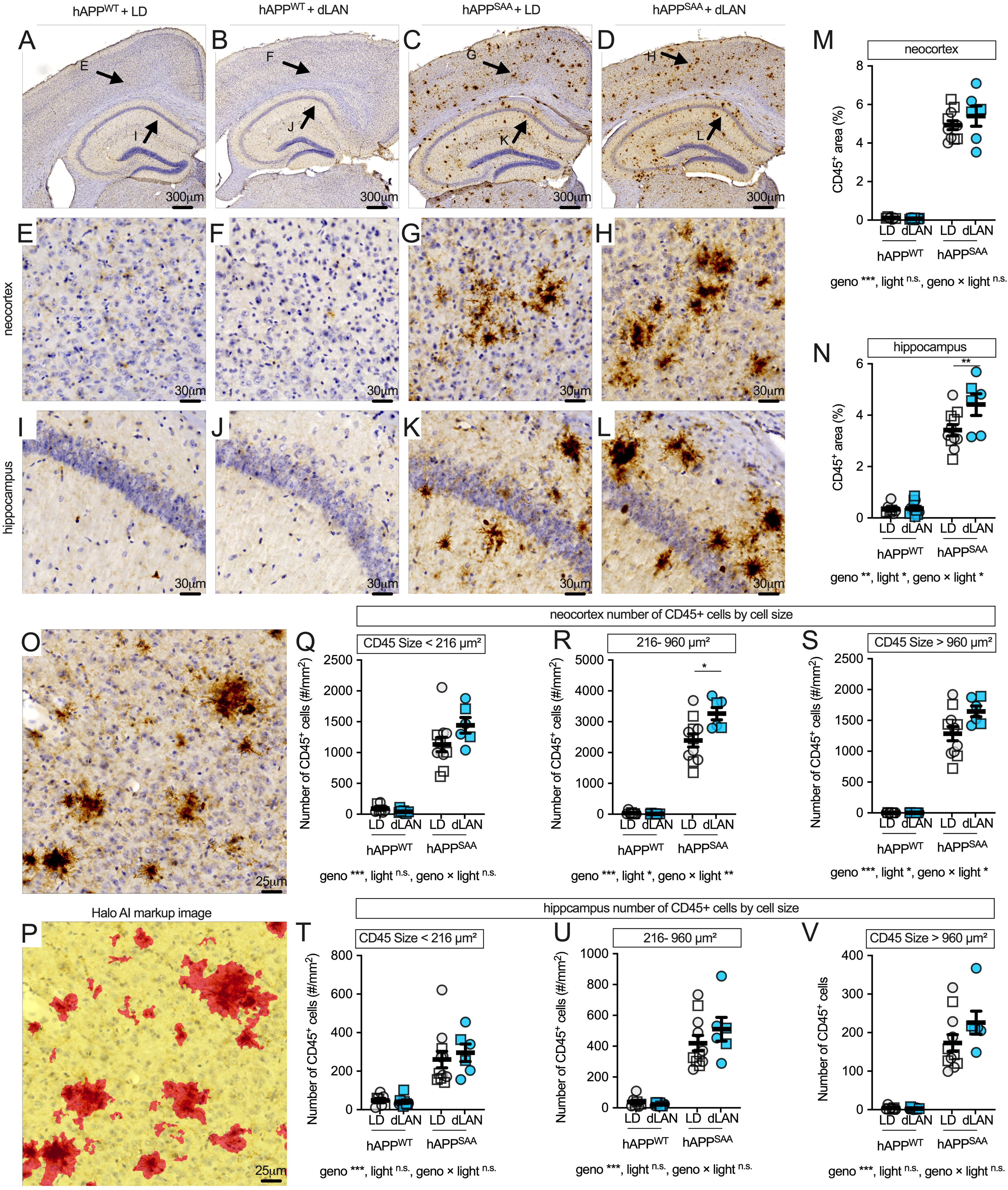
Dim light at night increases hippocampal CD45^+^ area in hAPP^SAA^ KI mice and shifts neocortical cluster-size distributions. (A-D) Low-magnification CD45 staining in hAPP^WT^ KI and hAPP^SAA^ KI mice maintained under a standard 12/12 light–dark cycle (LD) or dim light at night (dLAN). **(E-H)** Higher-magnification fields from neocortex; **(I-L)** hippocampus. **(M-N)** Quantification of CD45+ area (%) in neocortex and hippocampus. **(O)** Representative CD45+ clusters. **(P)** HALO AI markup of the same field as **O**, showing segmented CD45+ objects (red). Segmentation was used to define CD45+ clusters by size for quantitative analysis, with bins based on quartile-derived cut points: small, <216 µm² (≈16.6 µm diameter); medium, 216–956.5 µm² (≈16.6–34.9 µm diameter); and large, >956.5 µm² (≈34.9 µm diameter). **(Q–S)** Neocortical CD45+ cluster counts per mm² stratified by size (<227 µm²; 216–960 µm²; >960 µm²). **(T–V)** Hippocampal cluster counts per mm² stratified by the same bins. Symbols represent individual mice; circles denote females and squares denote males; bars show mean ± SEM. Sample sizes were: hAPP^WT^ KI-LD (n = 7, 4F/3M), hAPP^WT^ KI-dLAN (n = 12, 6F/6M), hAPP^SAA^ KI-LD (n = 12, 7F/5M), hAPP^SAA^ KI-dLAN (n = 8, 4F/4M). Analyses were conducted using two-way ANOVA (genotype × light) with planned within-genotype contrasts where appropriate. Main effects and interactions are indicated below the panels. Asterisks denote significance levels (*p < 0.05, **p < 0.01, ***p < 0.001).

Size-binned analyses of neocortical CD45+ clusters (HALO-AI segmentation; **Fig. 4O-P**) revealed robust genotype effects for small (<216 µm²), medium (216–960 µm²), and large (>960 µm²) clusters (all p < 0.001; **Fig. 4Q-S**). Light and genotype × light effects were significant for medium clusters (light: F(1,31) = 7.33, p = 0.011, partial η² = 0.191; interaction: F(1,31) = 8.27, p = 0.007, partial η² = 0.211) and for large clusters (light: F(1,31) = 5.00, p = 0.033, partial η² = 0.139; interaction: F(1,31) = 5.06, p = 0.032, partial η² = 0.140). Post hoc tests indicated an increase in medium-sized clusters in hAPP^SAA^ KI mice under dim light at night relative to the light–dark cycle (p = 0.023), with a trend toward more large clusters (p = 0.058). No statistical differences were seen with dim light at night exposure in hAPP^WT^ KI mice. In hippocampus, cluster counts per mm² were increased by genotype for all size bins (small: F(1,31) = 50.71; medium: F(1,31) = 102.76; large: F(1,31) = 121.87; all p < 0.001) and were not affected by light condition or interaction (all p ≥ 0.14; **Fig. 4T-V**). Dim light at night increased hippocampal CD45+ area in hAPP^SAA^ KI mice and modestly shifted neocortical CD45+ cluster-size distributions toward medium and large clusters, without altering hippocampal cluster counts. These changes mirror plaque pathology, suggesting that chronic light exposure at night may affect microglial plaque compaction. The divergence between area and number measures across regions suggests a potential transition point in pathological progression, which will require temporal profiling in future studies.

### Dim light at night increases MHCII^+^ cell counts in hAPP^SAA^ KI mice without consistent effects on hippocampal MHCII area

MHCII, a key antigen-presenting molecule, is induced by pro-inflammatory cytokines such as IFNγ and is often used as a marker of primed microglia. Expression is also found on infiltrating macrophages. MHCII immunolabeling was sparse in hAPP^WT^ KI mice and concentrated in clusters in hAPP^SAA^ KI mice (**Fig. 5A–F**). In contrast to CD45+, which appeared predominantly in clusters adjacent to plaques, MHCII labeling showed a strikingly different pattern, with regional localization, particularly in the hippocampus, that did not directly overlap with plaque distribution (**Fig. 5A-F**). In neocortex, MHCII+ area (%) was strongly increased by genotype (F(1,35) = 124.63, p < 0.001, partial η² = 0.781), while effects of light condition (F(1,35) = 3.05, p = 0.09) and genotype × light interaction (F(1,35) = 2.92, p = 0.096) did not reach significance. Similarly,MHCII+ area in hippocamus was also elevated by genotype (F(1,31) = 30.25, p < 0.001, partial η² = 0.494) but unchanged by light condition or interaction (**Fig. 5H**).

**Figure 5.**
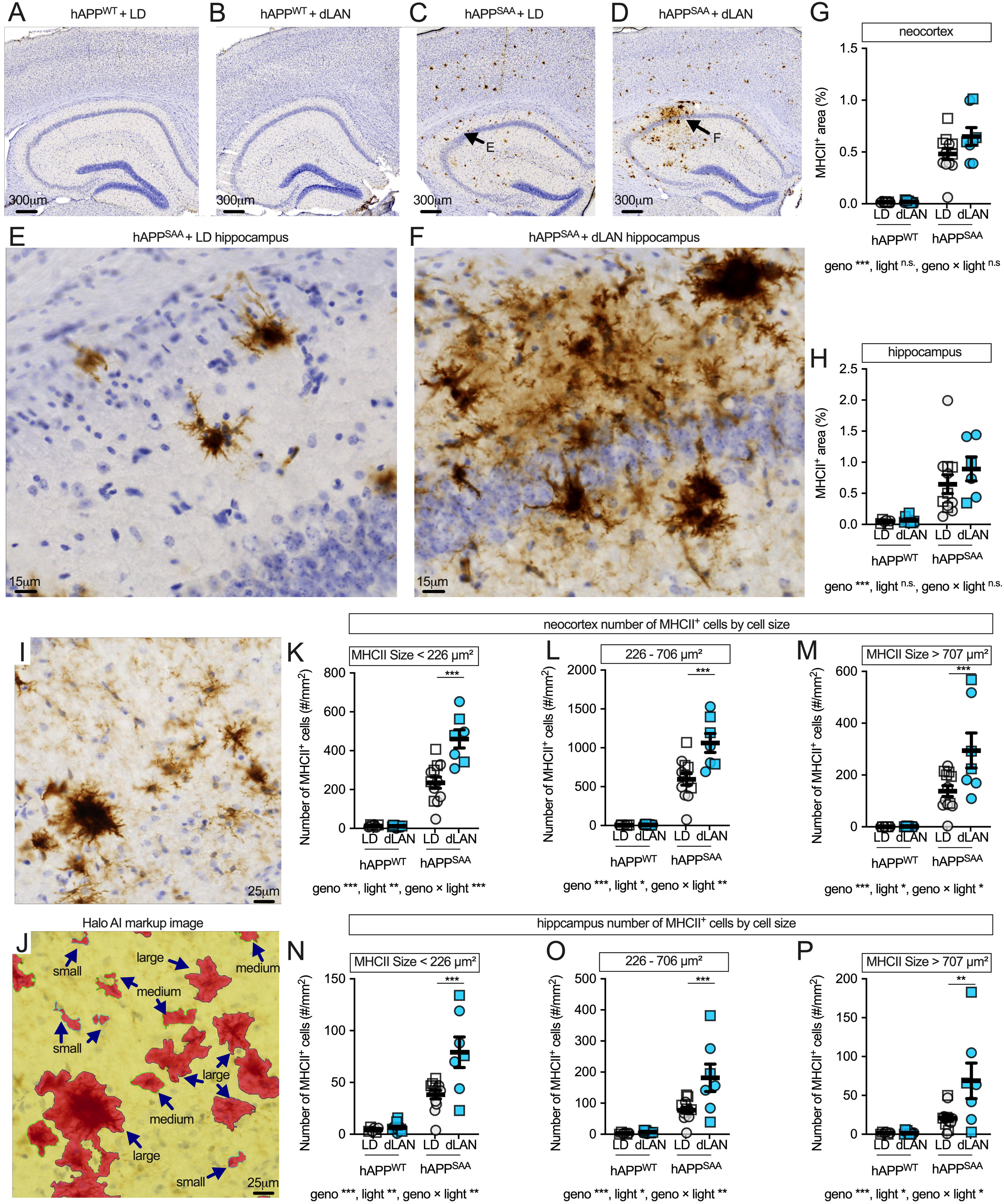
Dim light at night increases MHCII^+^ cell counts in hAPP^SAA^ KI mice. (A-D) Low-magnification MHCII staining in coronal sections from hAPP^WT^ KI and hAPP^SAA^ KI mice maintained under a standard 12/12 light–dark (LD) cycle or dim light at night (dLAN). **(E-F)** Higher-magnification neocortical fields showing clusters of MHCII+ cells (brown DAB; hematoxylin counterstain). Quantification of MHCII+ area (%) in neocortex **(G)** and hippocampus **(H)**. **(I)** Higher-magnification hippocampal field showing MHCII+ clusters. **(J)** HALO AI markup of the same region, illustrating automated segmentation of MHCII+ objects (red). Segmentation was used to define MHCII+ objects by size for quantitative analysis, with bins determined from quartile-based cut points of object area: small <226 µm² (≈ <16.9 µm diameter), medium 226– 706 µm² (≈ 16.9–30.0 µm diameter), and large >707 µm² (≈ >30.0 µm diameter). **(K-M)** Neocortical MHCII+ object counts per mm² stratified by size. **(N-P)** Hippocampal MHCII+ object counts using the same bins. Symbols represent individual mice; circles denote females and squares denote males; bars show mean ± SEM. Sample sizes were: hAPP^WT^ KI-LD (n = 7, 4F/3M), hAPP^WT^ KI-dLAN (n = 12, 6F/6M), hAPP^SAA^ KI-LD (n = 12, 7F/5M), hAPP^SAA^ KI-dLAN (n = 8, 4F/4M). Analyses were conducted using two-way ANOVA (genotype × light) with planned within-genotype contrasts where appropriate. Main effects and interactions are indicated below the panels. Asterisks denote significance levels (*p < 0.05, **p < 0.01, ***p < 0.001).

In neocortex, object-based analyses with HALO-AI segmentation (**Fig. 5I-J**) showed robust genotype effects across all object sizes. Small MHCII+ objects (<226 µm²) were increased by genotype (F(1,32) = 144.89, p < 0.001) and also showed effects of light (F(1,32) = 15.95, p < 0.001) and genotype × light interaction (F(1,32) = 16.59, p < 0.001). Post hoc contrasts confirmed more small objects in hAPP^SAA^ KI mice under dim light at night compared with standard light–dark cycle (p < 0.0001), with no effect in hAPP^WT^ KI mice. Medium objects (226– 706 µm²) were elevated by genotype (F(1,32) = 131.71, p < 0.001), with additional effects of light (F(1,32) = 10.85, p = 0.002) and genotype × light interaction (F(1,32) = 10.58, p = 0.003). Post hoc contrasts showed higher medium object counts in hAPP^SAA^ KI mice exposed to dim light at night compared with standard light– dark cycle (p < 0.0001), with no effect in hAPP^WT^ KI. Large objects (>707 µm²) were increased by genotype (F(1,32) = 48.88, p < 0.001), with significant effects of light (F(1,32) = 6.55, p = 0.015) and genotype × light interaction (F(1,32) = 6.34, p = 0.017). Post hoc tests indicated more large objects in hAPP^SAA^ KI mice under dim light at night compared with standard light–dark cycle (p < 0.001), while hAPP^WT^ KI showed no effect of light (**Fig. 5K-M**).

In hippocampus, object-based analyses showed strong genotype effects across all object sizes (**Fig. 5N-P**). Small objects (<226 µm²) were increased by genotype (F(1,32) = 65.15, p < 0.001), with significant effects of light (F(1,32) = 10.44, p = 0.003) and genotype × light interaction (F(1,32) = 8.69, p = 0.006). Post hoc contrasts confirmed more small objects in hAPP^SAA^ KI mice exposed to dim light at night compared with standard light–dark cycle (p < 0.0001), with no effect in hAPP^WT^ KI mice (p = 0.97) (**Fig. 5N**). Medium objects (226–706 µm²) were elevated by genotype (F(1,32) = 44.14, p < 0.001) and also showed significant light (F(1,32) = 8.29, p = 0.007) and genotype × light effects (F(1,32) = 7.16, p = 0.012). Post hoc contrasts again showed higher medium object counts in hAPP^SAA^ KI mice exposed to dim light at night compared with standard light–dark cycle (p < 0.0001), with no effect in hAPP^WT^ KI mice (p = 0.91) (**Fig. 5O**). Large objects (>707 µm²) were elevated by genotype (F(1,32) = 20.37, p < 0.001), with additional effects of light (F(1,32) = 6.25, p = 0.018) and genotype × light interaction (F(1,32) = 6.27, p = 0.018). Post hoc tests indicated more large objects in hAPP^SAA^ KI mice under dim light at night compared with standard light–dark cycle (p < 0.001), while hAPP^WT^ KI mice showed no light effect (**Fig. 5P**).

Together, these results indicate that dim light at night primes microglia toward an MHCII+ state, with synergistic effects of Aβ and light exposure evident only in hAPP^SAA^ KI mice. In contrast to CD45, which was predominantly plaque-associated, MHCII labeling did not map specifically to areas of plaques and showed broader regional localization, particularly in hippocampus. These patterns suggest that light exposure enhances MHCII-associated microglial activation outside plaque cores; whether this priming alters microglial functions relevant to plaque growth and compaction will require temporal and spatial profiling in future studies.

### Cortical and hippocampal cytokines and chemokines are elevated in hAPP^SAA^ KI mice, with minimal effects of dim light at night

Multiplex immunoassays of neocortical and hippocampal homogenates showed broad increases in inflammatory mediators in hAPP^SAA^ KI compared with hAPP^WT^ KI mice. In neocortex, genotype robustly increased IL-1β, IL-6, TNFα, and IL-17A (all p ≤ 0.003; partial η² = 0.23-0.79), whereas IL-10 was unchanged (p = 0.83) (**Fig. 6A**). Chemokines CCL2, CCL3, CXCL1, CXCL2, and CXCL10 were likewise elevated by genotype (all p < 0.001; partial η² = 0.64-0.87) (**Fig. 6B**). In hippocampus, genotype effects were again present for all cytokines except IL-6 (p = 0.24) (**Fig. 6C**) and for all chemokines (**Fig. 6D**) (Fig. 6C-D; p ≤ 0.02; partial η² = 0.15-0.90). Across analytes, light condition (light-dark vs dim light at night) and genotype × light interactions were not significant. The absence of a robust cytokine/chemokine signal at week 8 may reflect timing rather than insensitivity to light. Given that circadian disruption was most pronounced at week 2, inflammatory mediators may peak early and then normalize in parallel with partial recovery of circadian metrics, while microglial markers such as MHCII and CD45 fail to resolve. This raises the possibility that cytokine/chemokine induction is an early, transient event, whereas microglial activation reflects a failure of resolution. Future studies should test this by performing temporal cytokine profiling within the first 1-4 weeks of dim light at night exposure, and by blocking cytokine signaling during this window, to determine whether early inflammation drives the persistent MHCII/CD45 and Aβ changes.

**Figure 6.**
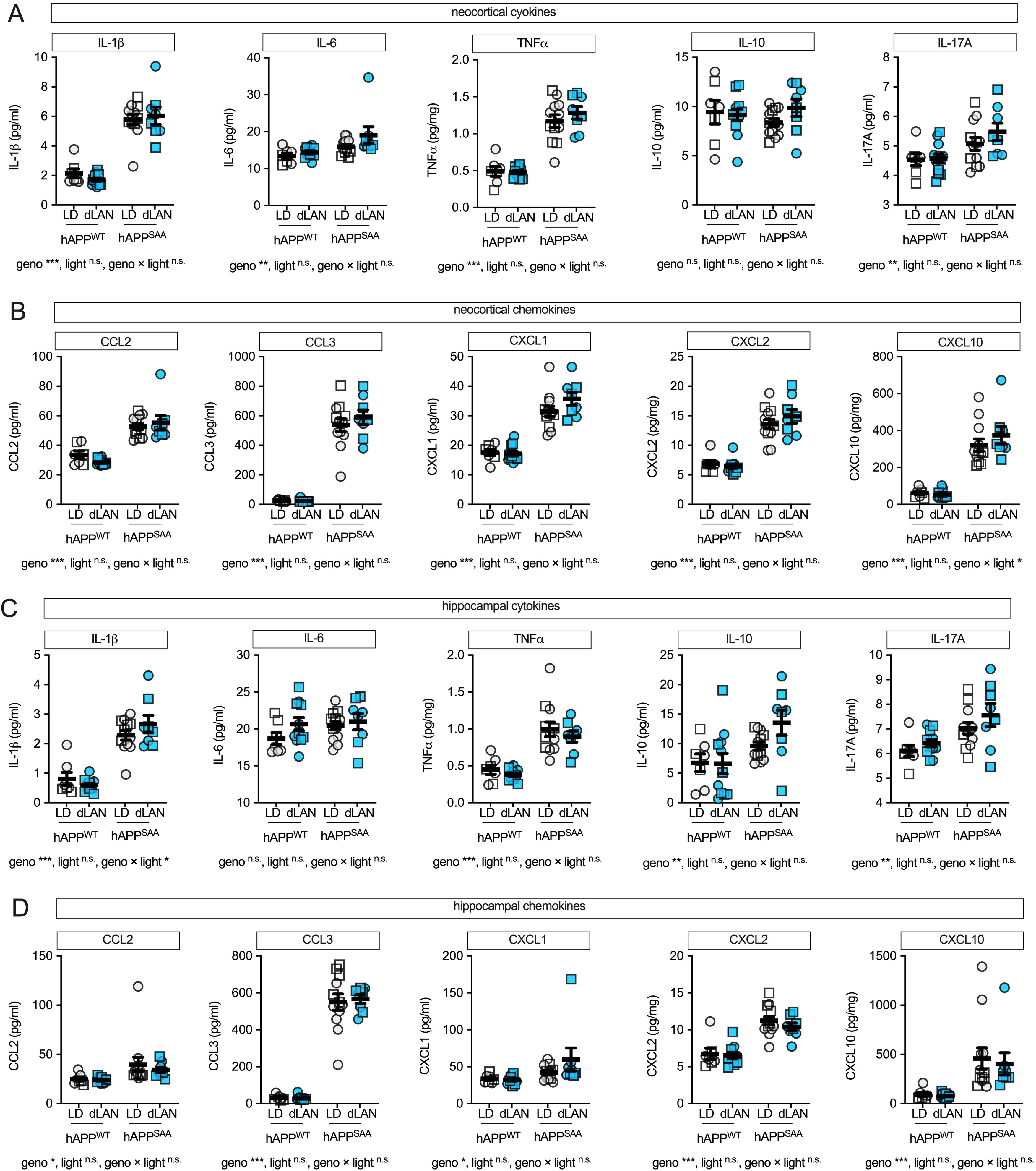
Pro-inflammatory cytokines and chemokines are elevated by amyloid genotype, with little additional effect of dim light at night. **(A)** Neocortical cytokines (IL-1β, IL-6, TNFα, IL-10, IL-17A). **(B)** Neocortical chemokines (CCL2, CCL3, CXCL1, CXCL2, CXCL10). **(C)** Hippocampal cytokines. **(D)** Hippocampal chemokines. Values are PBS-soluble cytokine and chemokine concentrations in tissue homogenates (pg/mg protein) measured using the MSD V-PLEX multiplex assay. Symbols represent individual mice; circles denote females and squares denote males; bars show mean ± SEM. Sample sizes were: hAPP^WT^ KI-12/12 light–dark (LD) (n = 7, 4F/3M), hAPP^WT^ KI-dim light at night (n = 12, 6F/6M), hAPP^SAA^ KI-LD (n = 12, 7F/5M), hAPP^SAA^ KI-dim light at night (n = 8, 4F/4M). Analyses were conducted using two-way ANOVA (genotype × light) with planned within-genotype contrasts where appropriate. Main effects and interactions are indicated below the panels. Asterisks denote significance levels (*p < 0.05, **p < 0.01, ***p < 0.001).

## Discussion

Light at night, also called light pollution, is increasingly recognized as an environmental factor that may influence cognitive decline ^25–27^. In the present study, chronic exposure of aged, humanized APP-KI mice to dim light at night (8 lux during the dark phase) disrupted circadian activity rhythms and altered neuropathological outcomes in an amyloid-prone background. We observed disruption within two weeks, with reduced amplitude and stability of the daily activity–rest rhythm and increased fragmentation. The pattern of reduced amplitude and increased fragmentation we observed has also been linked to higher AD incidence and faster progression from mild cognitive impairment in older adults ^10–12^. Our findings are consistent with prior work in other mouse strains showing that dim light at night reduces the amplitude of rhythms in food consumption, body temperature, and heart rate ^28^ and exacerbates microglial activation and neuroinflammation in the hippocampus after brain injury ^29,30^. For example, dim light exposure following experimental stroke or global ischemia increased proinflammatory cytokines including TNFα, IL-6, and IL-1β ^31,32^. Together, these findings suggest that dim light at night impairs circadian organization and, through altered microglial responses, may worsen neuropathology in vulnerable contexts such as amyloid deposition.

The association in humans between dim light at night exposure and AD ^27^ led us to investigate whether dim light at night exposure stimulates processes involved in AD progression, specifically amyloidosis and neuroinflammation. To this end we used two newly developed APP knock-in lines. The hAPP^SAA^ KI line carries a humanized Aβ region (G676R, F681Y, R684H) together with the KM670/671NL (Swedish), E693G (Arctic), and T714I (Austrian) mutations in the App gene, which drive amyloidogenic processing and result in robust plaque pathology in vivo without altering endogenous gene regulation. In contrast, the hAPP^WT^ KI line contains only a humanized Aβ sequence without disease-linked mutations. Dim light at night modestly increased hippocampal plaque burden and elevated soluble Aβ in neocortex of hAPP^SAA^ KI mice but had no effect in hAPP^WT^ KI mice. This distinction indicates that light exposure does not independently induce Aβ aggregation but rather exacerbates pathology in the context of a mutation-prone background, making hAPP^SAA^ KI a sensitized model for probing interactions between circadian disruption and amyloid disease mechanisms.

Astrocyte reactivity, as measured by GFAP, was markedly elevated in hAPP^SAA^ KI mice but was not altered by dim light at night. This lack of effect contrasts with prior studies showing that astrocytes are sensitive to sleep fragmentation and intermittent hypoxia, where GFAP increases were robust ^13,21^. To date, astrocyte responses to dim light at night have not been extensively evaluated, leaving it unclear whether the absence of an effect here reflects a true insensitivity to light-induced circadian disruption or simply a gap in the literature. These differences suggest that not all forms of circadian or sleep disruption exert equivalent effects on astrocytes. Whereas sleep fragmentation and hypoxia appear sufficient to drive astrocytic reactivity, light-induced circadian fragmentation did not further increase GFAP in the context of amyloid pathology, pointing instead to microglia as the more responsive glial population under these conditions.

Dim light at night altered microglial phenotype in ways that may have consequences for plaque dynamics. The shift toward larger CD45+ clusters and the increase in MHCII+ microglia observed only in hAPP^SAA^ KI mice suggest that light exposure does not simply amplify amyloid-associated microgliosis but changes its character. Whereas CD45 remained tightly plaque-associated, MHCII was more diffusely distributed, raising the possibility that dim light at night diverts microglia from compacting plaques towards a primed or antigen-presenting state. Prior work has shown that microglia contribute to plaque compaction, and that disruption of this function results in more diffuse and potentially more neurotoxic plaques ^33^. Thus, the shift toward an MHCII+ primed state may indicate that light exposure alters microglial involvement in plaque remodeling, favoring immune activation over compaction. These findings suggest that dim light at night modifies the quality of microglial engagement with plaques, potentially reducing their capacity to compact Aβ while diverting resources toward antigen presentation. Testing this possibility will require mechanistic experiments that directly probe microglial contributions to plaque morphology. Future studies could measure plaque density and morphology across timepoints to assess whether changes in microglial phenotype correspond to structural alterations, and could use targeted manipulations such as microglia-specific deletion of MHCII, blockade of IFNγ signaling, or modulation of TREM2 to determine whether the shift toward antigen presentation is causal. Pharmacological depletion and repopulation of microglia could further test whether microglial responsiveness to dim light at night is required for changes in plaque burden and organization. Such approaches would clarify whether circadian disruption diverts microglia away from a plaque-compacting role and toward sustained antigen presentation, thereby altering the trajectory of amyloid pathology.

Cytokine analyses showed broad increases in inflammatory mediators in hAPP^SAA^ KI mice, while light exposure had little additional effect. The absence of a cytokine response is notable, given that IL-1 and TNFα ^34,35^ are sleep-regulatory cytokines and are elevated by sleep fragmentation ^13^. In contrast, four weeks of exposure to dim light at night has been shown to exaggerate CNS IL-1β, TNFα, and IL-6 responses to LPS ^36^, high-fat diet ^15^, and particulate matter ^16^. There is also evidence that this exaggerated cytokine response is sex dependent, with females showing a greater effect ^37^. In our study, we did not detect evidence of cytokine priming despite the presence of Aβ pathology, which itself could be considered a chronic challenge. This difference suggests that dim light at night may influence inflammatory responses differently depending on whether the stimulus is acute or chronic. It also remains possible that cytokine changes occurred early and normalized by the time of sampling, leaving shifts in microglial phenotype as the prevailing consequence of light exposure. Consistent with this idea, four weeks of dim blue light (∼5 lux) at night increased microglial number, altered morphology, and elevated iNOS even without immune stimulation ^29^. Taken together, these results suggest that dim light at night can alter inflammatory tone, but in the setting of amyloid pathology, the more persistent effect may be failure of microglial resolution rather than sustained cytokine production.

Several limitations should be considered. Pathological analyses were performed at a single timepoint, limiting conclusions about the temporal progression of immune and amyloid changes. The results of the current study did not reveal significant interactions between dim light at night and biological sex on the parameters investigated. Prior work has reported sex-related differences in sleep, Aβ pathology, and inflammatory responses, including differential susceptibility to sleep fragmentation ^3^. Thus, the absence of sex effects here should be interpreted cautiously, as the present study was not powered to detect sex-specific interactions. Effect size estimates from the current dataset can inform the design of future studies specifically powered to determine whether sex modifies the impact of dim light at night on circadian function. In addition, cognitive behavior and synaptic physiology were not assessed, so the functional consequences of dim light at night exposure remain uncertain. Finally, group sizes were not fully balanced, which may have reduced power for some comparisons.

In conclusion, dim light at night broadly disrupted circadian activity rhythms and increased markers of microglial activation in an amyloid-prone background. The persistence of CD45 and MHCII changes, even as circadian fragmentation partially normalized, suggests that early circadian disruption may initiate microglial responses that fail to resolve. This interaction between light exposure and amyloid pathology may represent one pathway through which chronic exposure to dim light at night influences neurodegeneration. The findings add to a growing body of evidence indicating that light at night has detrimental effects on numerous physiological and behavioral functions in humans ^4,6–8,38–41^ as well as in rodents ^15,16,31,32,36,42–49^. Together with prior work, these findings suggest that dim light at night is not only disruptive to circadian and metabolic functions but may also contribute to the progression of Alzheimer’s disease.

## Acknowledgments

Equipment used in this study was supported by the Office of the Vice President for Research at the University of Kentucky (SS and MJD). Research was supported by the National Institutes of Health (NIH R01 AG068215; MPM, MJD, ADB, BFO, and SS).

## Author Contributions

MJD and HRW designed the study. MPM, MJD, BFO, SS, and ADB provided funding and contributed to study design. ADB performed the statistical analyses and, together with MJD, prepared the initial draft. Experimental work was conducted by MRH, LB, HRW, SS, MFC, EGD, TM, KNR, KK, MCS, and CEJ under the supervision of senior authors. All authors contributed to data acquisition, analysis, or interpretation; critically revised the manuscript; approved the final version; and agree to be accountable for the integrity and accuracy of the work.

## Declaration of generative AI usage in scientific writing

During the preparation of this work, the author(s) used ChatGPT and Claude to assist in evaluating the original text for clarity, completeness, and style. These tools provided editorial suggestions and recommendations; however, the author(s) reviewed and edited all content as necessary. The author(s) take full responsibility for the final content of the published article.

